# SLiMAn: an integrative web server for exploring short linear motif-mediated interactions in interactomes

**DOI:** 10.1101/2022.01.14.476361

**Authors:** Victor Reys, Gilles Labesse

**Affiliations:** Centre de Biologie Structurale, Montpellier, France; La Ligue Contre le Cancer, Paris, France; Université de Montpellier, Montpellier, France

## Abstract

Cells express thousands of macromolecules, and their functioning relies on multiple networks of intermolecular interactions. These interactions can be experimentally determine at different spatial and temporal resolutions. But, physical interfaces are not often delineated directly especially in high-throughput experiments. However, numerous three-dimensional structures of complexes have been already solved and sequence conservation allows comparative modeling of additional complexes. A large fraction of protein-protein interactions involves domain and so-called SLiMs (for Short Linear Motifs). Often, SLiMs lie in disordered regions or loops. Their small size and loosely folded nature prevent straightforward detection. SLiMAn (Short Linear Motif Analysis), a new web server is provided to help thorough analysis of interactomics data. Starting from a list of putative interactants such as the output of an interactomics study, SLiMs (from ELM) and SLiM-recognition domains (from Pfam) are extracted and potential pairing are displayed. Additionally, filters are available to dig into the predicted results such as the motif E-value, IUpred2 scoring functions for disorder or BioGRID interaction matches. When structural templates are available, a given SLiM and its recognition domain can be modeled using SCWRL. We illustrate, here, the use of SLiMAn on three distinct examples including one real-case study. We oversee wide-range applications for SLiMAn in the context of massive analysis of protein-protein interactions at proteome-wide scale. This new web server is made freely available at http://sliman.cbs.cnrs.fr.

## Introduction

Proteins are essential cellular components involved in many functions including biosyntheses, structural roles and signal cascades. Many of these functions require multiple interactions. ^1,2^

Experimental interactomic studies are now unraveling potential partners at high pace and (sub-)proteomic level.^3,4^ Interactomic studies are usually performed using either genetic tools, such as two-hybrid system (Y2H), or a flagged protein serving as a bait to capture its interacting partners (e.g.: by pull down or immunoprecipitation), followed by an identification process (mass spectrometry, western-blot, … ). For example, HuRI ^3^ brings unprecedented information on 53,000 direct protein-protein interactions in *Homo sapiens* and encompasses 17,500 protein chains – over the 23,000 human proteins identified. Bottom-up mass spectrometry allows rapid detection of numerous preys for a given bait within a given cellular context, ^4^ but it does not provide clues for direct interactions and rather reveal macromolecular complexes or intricate networks of interactions.

Analyzing interactomic results aim to rebuild the protein-protein interaction (PPI) networks to unravel the mechanisms of biological processes, possibly at a molecular level. But the resulting interactomes are most frequently displayed as a raw list of identified proteins, annotated according to their biological function. ^5,6^ While already informative, these lists do not allow to draw the underlining macromolecular networks, nor to precise the structural interfaces involved. Solving all the corresponding 3D structures is currently not feasible. Hence, alternative approaches are required to validate or identify the actual interactions between each protein pair.

PPI network reconstruction remains a challenge, although some databases are devoted to help such a task. The STRING-db^7^ database, well appreciated by the community, combines genetic interactions, text mining, text association, predicted and experimental PPI databases to provide an overview of the corresponding networks. However, it usually lacks of structural information on the macromolecular interactions involved in the corresponding (sub-)networks although it indicates the knowledge (or not) of the structure for each partner. Similarly PPI databases, such as the Biological General Repository for Interaction Datasets (BioGRID^8^) gather the results of multiple interactomics studies, but are also lacking structural information.

Combining proteomics data with 3D search and comparative modeling has now emerged as a valuable tool to compensate limitations of each of the above approaches. Servers such as Interactome3D^9^ and Proteo3Dnet^10,11^ allow one to dig efficiently into available 3D structures, gathered in the Protein Data Bank (PDB^12^) to unravel likely physical interactions within a set of proteins. The Complex Portal (https://www.ebi.ac.uk/complexportal/) is also often used to match the proteins involved in 297 already known mammalian complexes at the structural level. However, they mainly focus on domain-domain interactions, while a vast number of PPIs involve so-called short linear motifs (SLiMs) in one partner and a recognition domain in another partner.^2^ They usually correspond to short segments (4-12 residues) and are frequently encountered in disordered regions or (long) loops although exception exist, while dedicated folded domains are involved in their specific recognition.^13^

The Eukaryotic Linear Motif resource (ELM^14^), is a reference database for SLiM annotation and prediction. From 3559 publications this database gathers 289 manually curated SLiM classes. Classes are defined by the signature – or patterns – of the binding motifs which are also associated with interacting and structured domains as annotated from the Protein Familly (Pfam) database.^15^ Each pattern is associated with an E-value estimating its frequency in protein sequences. The ELM database is now largely used by the community to identify potential SLiMs-mediated PPI. The main difficulty is the huge number of predicted interaction motifs for a given protein (over 200 per human protein on average), among which many are false positive matches.

More frequently found in intrinsically disordered regions of proteins or in external accessible loops, SLiMs can be considered to act as peptides that bind onto a target structured domain.^16^ For this reason, the current state-of-the-art for SLiM predictions also includes additional features, such as prediction of the disorder state of the SLiM. The software IUpred2^17^ is often used to predict the probability for a given amino-acid to be in a disordered part of a protein. Despite this efficient filtering, the analysis of a given data sets for an interactomic analysis is leading to a large number of matches. Furthermore, systematically filtering out SLiMs with high E-value (e.g.: E-value >0.005) or predicted as too ordered (IUpred <0.3) may exclude various SLiMs (e.g.: PDZ motifs which are often too close to a folded domain). Currently, performed on one protein at a time, this constitutes a huge burden when one is analyzing an interactomic output made of hundred(s) of proteins. Alternatively, performing such analysis on larger scale implies to fix various thresholds (E-value, IUpred2 scores, …). This is expected to limit the scope of tools such PrePPI^18^ which otherwise represents an interesting large-scale analysis of all potential SLiMs in the proteomes of important model organisms.

Here, we describe a new bioinformatic tool, SLiMAn dedicated to SLiMs analysis within sub-proteomes, based on an advanced selection scheme. This new tool allows focused annotation of protein motifs to discover truly interacting partners in a sub-proteome-wide manner.

The developed analysis tool is made freely accessible to the community through a dedicated server (http://sliman.cbs.cnrs.fr).

## Materials and methods

### Databases

#### Sequence annotation

SLiMAn relies on the subset of ‘reviewed’ entries from UniprotKB, ^19^ which corresponds to 564 277 entries (20 396 human), in the last update (July 2021). From each UniprotKB, Pfam annotations^15^ are extracted (domain boundaries and descriptions). From the ELM database,^14^ class names, interaction domain types (Pfam annotation), regular expression and E-value are extracted for SLiMs analysis. In the current release (July 2021), a total of 289 classes are defined. ELM experimental instances, ones that are used for the class regular expression definition, are also included, allowing fast validation for known interactions.

The PhosphoSitePlus®^20^ database, focusing on post-translational modifications (PTM) annotations, is integrated to pinpoint locations of acetylation, methylation, O-GalNAc, O-GlcNAc, phosphorylation, sumoylation and ubiquitination sites over the amino acid sequence for 46 096 proteins (from which 18 021 are human).

#### Protein-protein interaction data

The latest release (July 2021) of the BioGRID^8^ dataset was integrated in SLiMAn. Only physical interactions were retained, and split into low and high throughput experiments. The mapping between UniprotKB entry names and BioGIRD data is achieved using the Uniprot mapping API (https://www.uniprot.org/uploadlists/).

#### Structural information

For each of the possible 291 ELM/Pfam associations, the Protein Data Bank (PDB) ^12^ was parsed in search for structural information, using the pdb pfamA reg database to select the corresponding domain chains. For each referenced domain chain, other chains shorter than 35 residues in length, are converted to FASTA format, and associated ELM regular expressions are used to parse the sequences. During this conversion, modified residues (MLZ, MLY, M3L, ALY, SEP, TPR, MSE, MNN, DA2, SEC, TPO and PTR) found in the structure are converted to the one-letter code of the corresponding unmodified amino-acids (e.g.: S for SEP). To check actual peptide-domain interaction, all C*α* from residues that matched the regular expression must be found under a 10 Å threshold from any atoms of the domain chain. Then, contact distances between residues belonging to the peptide-domain interface are split into 4 categories (>.0 Å, 5.5-7 Å, 4.0-5.5 Å and <4.0 Å). These distance categories are converted into sequences of contact-scores (respectively 0, 1, 2 and 3), allowing a discretized encoding of the interface over the sequences of the domain and the peptide. After template extraction, domain amino-acid sequences are placed in a FASTA file and the BLAST^21^ database generator (*makeblastdb*) is used to setup the corresponding *blastable* database of the domains, for future alignment queries.

The parsing led to the extraction of 5064 templates (from 2228 distinct structures). The resulting database allows fast and precise template selection and SLiM-domain modeling for 201 ELM/Pfam associations for which at least one template was found. For the remaining 90 associations, no template could be extracted currently and therefore alignments/modeling can not be performed.

### Embedded software

#### Disorder prediction

IUpred2A^17^ is used to predict disorder along the amino-acid sequence of a protein. The disorder scores (ranging from 0 – most ordered – to 1 – most disordered) are predicted at the residue level and includes several predictors; local disorder (short and long window size corresponding to 25 and 100 residues, respectively), presence of structured domain (short and long windows) and ANCHOR2 (probability to be part of an interacting segment). The final scores attributed to the motif are obtained by averaging the scores over the residues constituting a given motif. An additional binary value is computed (StrictDisorder). It is set to 1 (0 otherwise) if all residues from a motif have their short, long and ANCHROR2 predictions above the 0.5 threshold.

#### Sequence alignments

Alignments between the domain sequence and template structure sequence is performed by two software. MAFFT^22^ is used with the local-pair alignment option, limited to 1000 max-iterations (L-INS-i). BLAST^21^ (*blastp*) is used with the pre-computed *blastable* databases, previously described.

For motif alignments, only MAFFT is used and the gap opening and extension penalty options have been increased to 10 and 0.2 respectively.

#### Alignments metrics

Five different alignment metrics are computed to guide selection of the most suitable templates for comparative modeling:

- Sequence Identity (%Ident), corresponding to the sum of identical aligned amino acids devided by the number of aligned amino-acids.
- Query coverage (%QueryCoverage), corresponds to the sum of aligned amino acids from the query divided by its length, and represents the percentage of the query amino-acids that will be modeled.
- Template coverage (%TemplateCoverage), corresponds to the sum of aligned amino acids from the template divided by the template length, and represents the percentage of the template amino-acids that will be used for the modeling. It is computed for both the ELM motif and the Pfam domain.
- Contact conservation score (CCS), corresponds to the sum contact-scores of aligned amino acids from the template. This metric (and its percent representation) allows fast discrimination of alignments that will lead to SLiM-domain interactions.

All the above described metrics should help better and faster selection of optimal sequence-structure alignments.

#### Comparative modeling

Three-dimension model of the SLiM-domain complex can be built from the sequence structure alignments. Here, only aligned and substituted residues have their side-chain conformations quickly optimized by SCWRL 3,^23^ with amino-acid backbone atoms and strictly conserved side-chains kept fixed. Modeling of the selected complex is then preformed in two steps. In the first step, the queried domain is modeled, using the extracted domain as template and the similarly extracted peptide as a constraint to model domain substituted amino-acids side-chains. Then, the resulting domain model is used as as a fixed constraint, to optimize side-chains of the interacting motif.

### Algorithm

Starting from a list of putative interactants (defined by their Uniprot accession numbers or entry names), SLiMAn analyzes the provided data at 3 successive levels, named SLiM-IP, SLiM-ID and SLiM-IM:

- First, all predicted ELM/Pfam pairing are displayed.
- Then, for each hit, the corresponding sequences can be aligned to related structural templates (if available).
- Finally, the resulting structural alignments can be submitted to a dedicated comparative modeling procedure and the resulting models can be directly vizualized (Fig. 1).

**Figure 1:**
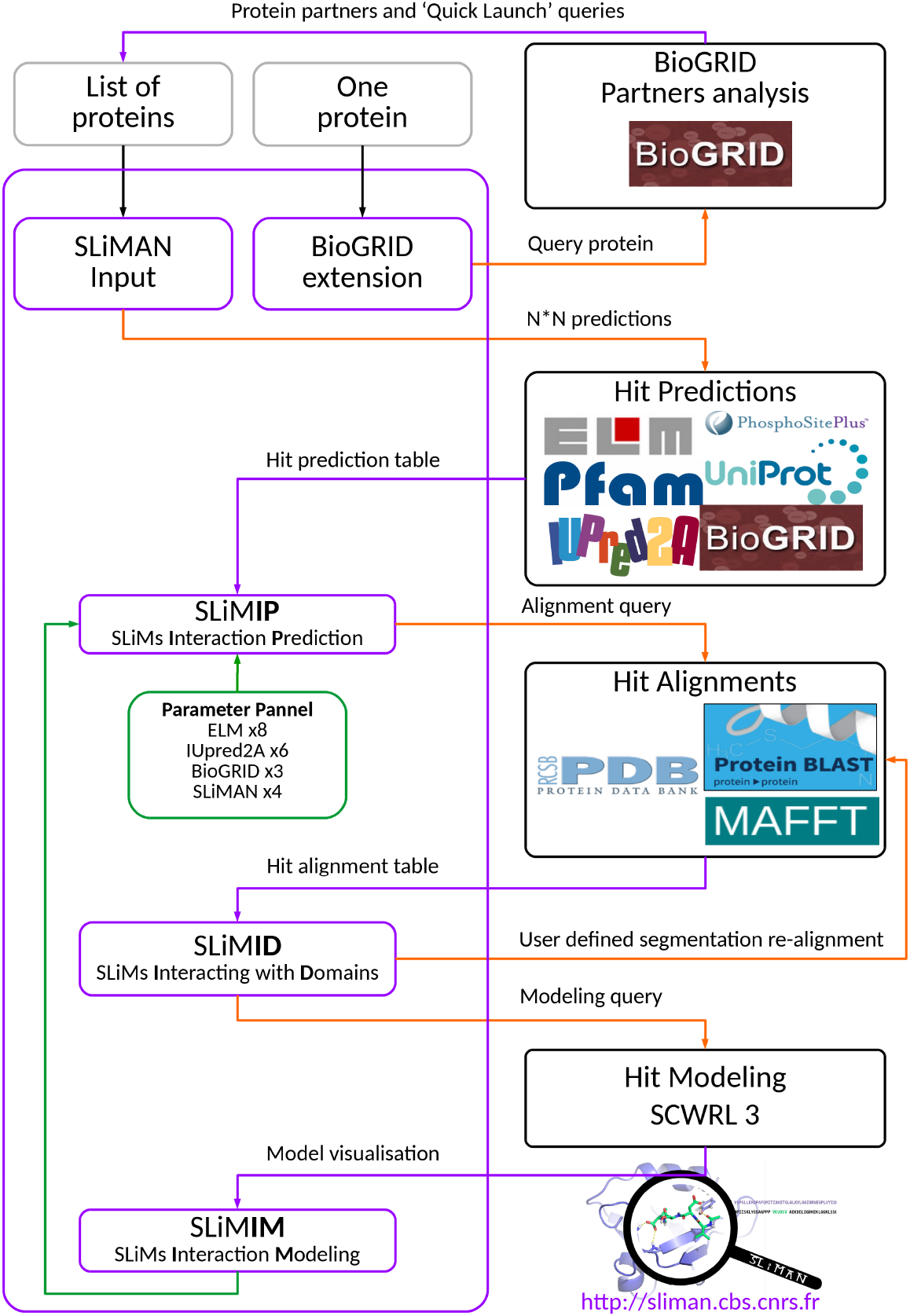
Detailed SLiMAn workflow from input putative partners to comparative modeling of the interactions. **SLiMAn one-sequence input:** this module searches for interacting partners in BioGRID. **BioGRID extension:** Interacting partners are listed and using ‘Quick Launch’, the whole list can be directly submitted to SLiMAn. **SLiMAn proteome input:** for a query list of putative interactants, matching entries from the ‘reviewed’ Unipro-tKB database are extracted for SLiM analysis. **Hit Predictions:** ELM motifs are paired with Pfam domains within the queried sub-proteome. **SLiM-IP:** hit predictions are displayed. Thresholds (IUPRED, E-value, …) can be adjusted. **Hit Alignment:** structural alignment for a given SLiM-domain interaction are provided when mtching templates are available. **SLiM-ID:** SLiM and Pfam domain are displayed, along with their alignments whose boundaries can be edited. Templates can be selected for comparative modeling. **Hit Modeling:** using the selected templates, the queried SLiM-domain interaction is modeled using SCWRL3. **SLiM-IM:** 3D models are listed and viewable using a JSmol applet. They can be downloaded. They can be filtered in or out, and this information fed back to SLiM-IP. or a list of UNIPROT names separated by a coma (“FRAT2 HUMAN, GSK3B HUMAN”). First, input entries are checked, and then the initialization of the new project is triggered to find motifs and matching domains.

The whole procedure is performed in a stepwise and interactive manner.

#### Hit predictions

Regular expressions from the linear motifs referenced in ELM are used to parse the sequences of the input proteins. For each regular expression detected, all Pfam domains found in the input proteins and matching the ELM class are also extracted. If no Pfam domain matches a given ELM motif, the latter is dropped out. For each motif, IUpred2A is used to compute the average disorder scores (Short, Long, ShortDom, LongDom, Anchor2 and StrictDisorder). In addition, for the two partners paired, the PPI database (BioGRID) is searched and high and low throughput experimental interactions are counted. The final prediction, for a given motif-domain association (hit), is a set of 17 descriptors (see Supplementary Table 1). The four most important descriptors (ELM E-Value, IUpred StrictDisorder and BioGRID LowThrougput and Total counts) are used to compute a confidence score. All predicted associations (hits) are written in a tabular separated value file.

Hit predictions are displayed in the SLiM-IP (SLiMs Interaction Prediction) page, as a table with ELM motifs in the left columns and interacting domains in rows. Sequence segments corresponding to an ELM motifs are displayed with the corresponding PTM from PhosphoSitePlus® annotation, when available.

On the same page, a interactive parameter panel is displayed with default parameters (see Supplementary Table 2). Users can modify a set of 21 different parameters, and navigate into the predicted results with distinct stringency. Additionally, three summary tables are displayed (see Supplementary Table 3) to recapitulate the number of ELM motifs and Pfam domains found and filtered (in/out), the ELM classes found and filtered as well the total number of interactions predicted.

Displayed hits are colorized according to the confidence score (from 1 to 4), and links to alignments and models are displayed when templates are available. By clicking on the ‘Alignments’ link, a alignment query of the corresponding hit is launched.

#### Hit alignments

For a selected hit, motif and domain sequences are aligned (c.f. Sequence Alignments ) with templates corresponding to the same ELM-Pfam combination.

Alignments results are displayed on the SLiM-ID (SLiMs Interacting with Domains) page. Matching residues from the motif and the domain are highlighted (in green) on the full-length sequences. Domain/motif boundaries can be modified at will. Paired sequence-structure alignments are displayed in a dedicated table with a color code indicating either the conservation (query sequences) and the contacts (template sequences) for both the domain and the peptide motif. Alignments can be sorted by the different alignment metrics.

Users can select and submit the desired pairs of motif/domain alignments for comparative modeling.

#### Interaction modeling

Quick modeling is based on optimization of side-chain conformations within the Pfam domain and then the ELM motif using SCWRL3 (c.f. Comparative Modeling). Generated models are displayed in the SLiM-IM section (SLiMs Interaction Modeling). Models are accessible in a table and can be downloaded or visualized online using the JSmol applet (https://sourceforge.net/projects/jsmol^24^). Users are allowed to select in/out promising/incorrect modeles of complexes, and the corresponding information is brought back into the hit prediction table in SLiM-IP. This may help further analysis of the protein network.

### Implementation

SLiMAn (http://sliman.cbs.cnrs.fr) was designed as a web application, enabling a visual and interactive representation of the potential interactions involving SLiMs.

On the SLiMAn homepage, users are asked to input a list of proteins, to start a new SLiMAn project (see: http://sliman.cbs.cnrs.fr/Documentation.html). The inputs should be formatted either as a list of FASTA headers (e.g.: “ sp∥O75474∥FRAT2 HUMAN”)

#### BioGRID extension

Another feature proposed by SLiMAn in the input section is the BioGRID extension analysis. On this extension, the BioGRID database is analyzed with the input protein(s), and the resulting interactants (for each entry) are displayed in a table, sorted by number of interactions. This list can, in turn, be submitted to SLiMAn using the ‘QuickLaunch’ button (or by copy-paste of – part of – the list of interactants to the SLiMAn input section).

#### Server

SLiMAn runs on a virtual machine with 4 CPU and 8Go of RAM under Ubuntu 18.04 environment. Apache2.0 is used as web server. Python3 and R programming languages are used for computations while rendering is done using HTML, CSS, JavaScript and CGI scripts. Implemented databases are stored in CSV or JSON format files. Computed data are stored in conventional text formats (CSV, JSON, PDB and FASTA), and always available for downloads. Supplementary documentation, source code and test files are available at : http://sliman.cbs.cnrs.fr/Documentation.html.

#### Datasets

The first example was build by gathering B-Raf with some 14-3-3 partners. The dataset for FRAT2 was extracted from BioGRID and contained 13 partners. The Pragmin data set was retrieved from the literature^25^ and corresponds to a SILAC-based quantitative proteomic analysis, which allowed the identification in a semi-quantitative manner of Pragmin interactors in FLAG-Pragmin transfected human HEK293T cells. More detailed information can be found in the original paper.

## Results and discussion

Below, we describe in some details three distinct examples. The first one illustrated the use of SLiMAn to extract from the PDB templates for comparative structure modeling that perfectly match a given ELM/Pfam pair. The second example deals with exploration meta-interactomics data from BioGRID to unravel new interaction. Serendipitously, the proposed interaction matches one involving a close homologue that was validated experimentally.^26^ The last example corresponds to a real-case study of an interactome recently described,^27^ in an exploratory mode. Some other pre-computed examples have been also made available on the index page of this web server.

### B-Raf / 14-3-3

A simple example indicates the role of parameter tuning to unravel some motifs and the interest of the modeling module in SLiMAn. The protein-kinase B-Raf has been shown to interact with 14-3-3 proteins. 14-3-3 recognition motif have a rather high E-value of 0.004477. This suggests high chance of finding spurious motifs. However, SLiMAn showed that two 14-3-3 binding motif of B-Raf (residues 362-367 and 437-442) occur in a largely unfolded segments (STRICT DISORDER: YES). Furthermore, the phosphorylation of a serine/threonine in the motif is documented in PhosphoSitePlus®. The structure of B-Raf motif bound to a human 14-3-3 protein can be modeled easily using the interface with SCWRL and the experimental structure corresponds to an hybrid human/insect complex. ^28^ Of note, the phosphorylation is not modeled so far.

### FRAT2 network

To illustrate the use and power of the new tool, we describe, below, a short and clear example with an experimental validation provided from the literature. FRAT2 is an inhibitor of GSK3*β*, which is a well-studied protein-kinase known to shuttle between the nucleus and the cytoplasm. ^26^ Interestingly, interrogating usual databases like Uniprot, PubMed, BioGRID or STRING-db does not provide any indication for the mechanism involved in this translocation.

To find a possible mechanism of translocation, we used SLiMAn to extract interacting partners of human FRAT2 from BioGRID. At the time the query was made (July 2021), a total of 13 interacting partners of FRAT2 were retrieved, including GSK3*β* and XPO1. Using default parameters (as listed in Supplementary Table 2), SLiMAn highlights 12 possible interactions (see http://sliman.cbs.cnrs.fr/FRAT2_DEFAULT) based on 9 SLiMs found in 3 proteins (FRAT2, GSK3*β* and BUB1). Those SLiMs matched with 2 types of Pfam domains (PF00069 and PF08389) detected in 3 proteins (XPO1, GSK3*β* and BUB1). Of note, motifs corresponding to phosphorylation (MOD class in ELM) are precisely associated to a given sub-family of protein-kinases (NEK2, PKB or CDK in this example) while they are associated to the same Pfam domain PF00069 corresponding to most of known protein-kinases. Hence, most association are spurious. Here, they would correspond to auto-phosphorylation of GSK3*β* or BUB1. Nevertheless, the 3 motifs highlighted, here, do correspond to phosphorylation sites (according to PhosphoSitePlus®), but are likely due to other protein-kinases while BioGRID associations would be due to homodimerization of those two protein-kinases. In parallel, 8 motifs from the DOC class were also detected but again with an annotated specificity (MAPK) suggesting they might not be relevant here.

Using SLiMAn, within a few clicks, we were able to identify and model a Nuclear Export Signal (NES) motif present in FRAT2 (Fig. 2). GSK3*β* and FRAT2 have no documented NES motif in Uniprot, but SLiMAn straightforwardly highlights that FRAT2 harbors such a motif. FRAT2 has been found to interact with a major exportin XPO1 in a high-throughput experiment.^29^ The motif shows favorable sequence parameters (E-value = 0.0007626; IUpred short score= 0.42 and ANCHOR2 = 0.809). Note that FRAT2 is predicted to be mainly natively unfolded beside short helical segments. Interestingly, an experimental validation using directed mutagenesis of the NES signal in the homologous FRAT1 was performed almost a decade ago^26^ but this information did not make its way to most databases. Satisfactorily, the NES motif highlighted in FRAT2 by SLiMAn matched the validated NES in FRAT1. To gain further insight, we used SLiMAn for homology modeling of the corresponding interface with XPO1 (Fig. 2) after slight adjustment of the sequence-structure alignments. By default, the PF08389 domain of XPO1 supposedly runs from residues 123 to 268, and the domain and the motif stand far apart in the corresponding model. In fact, the boundaries of this long ARM-repeats should be extended to include the actual NES binding site (centered around position 571). To focus on the interacting region, we set the XPO1 domain to run from residues 520 to 620 in SLiM-ID, and resume the modeling task (Fig. 2). Careful analysis of the contacting residues at the motif-domain interface suggested a one-residue shift to optimize the interaction and especially to bury some hydrophobic residues of FRAT2 (L51 and L53) within the binding groove of XPO1 (as observed in other related complexes). To obtain a better alignment of the motif with its templates, motif boundaries were changed from 41-54 to 42-53. These changes led to a model showing favorable interactions all along the 12-residue long motif that contains a short helix and an extended segment (see http://sliman.cbs.cnrs.fr/FRAT2_HUMAN). Noteworthy, mutation to alanine of two leucines (corresponding to L51 and L53 in FRAT2) in FRAT1 abrogates GSK3*β* nucleus export.^26^ Our modeling also reveals that the binding occurs close to the E571 residue in XPO1 that is mutated in chronic lymphocytic leukemia.^30^ Accordingly, not only a clearer picture of the shuttling mechanism of GSK3*β* was unraveled but also a new hypothesis could be drawn for the role of the mutation of XPO1 in a particular cancer.

**Figure 2:**
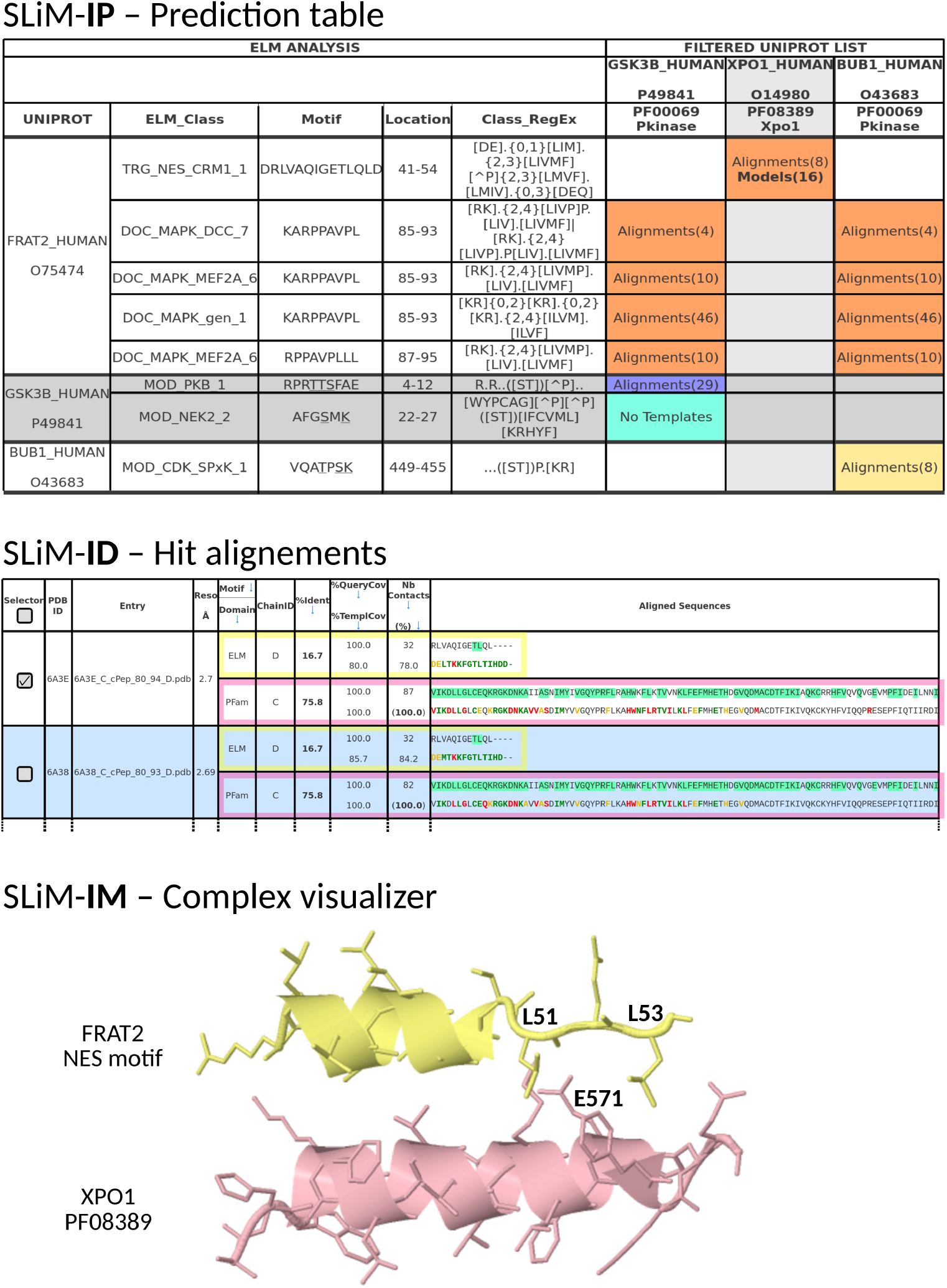
SLiMAn outputs focused on FRAT2 interactions, analysed by SLiM-IP, SLiM-ID and SLiM-IM. **SLiM-IP:** Table of potentially interacting pairs of motifs/domains, colored by their confidence score. **SLiM-ID:** Table holding sequence-structure alignments for the FRAT2 NES motif and XPO1. On each template rows, both motif and domain sequence-structure alignments are displayed. Query sequences are highlighted according to sequence Identity. Template sequences are colored according to the contacts scores (black, red, yellow, and green for respectively >7.0 Å, 5.5-7 Å, 4.0-5.5 Å and <4.0 Å). **SLiM-IM:** Comparative modeling, from PDB 6A3E template, of the FRAT2 NES motif (yellow) in complex with XPO1 (pink), visualized online by the JSmol applet.

Next, we compared our results with the outputs from other tools dedicated to provide structural insight into protein networks (Fig. 3). For example, the recently developed Proteo3Dnet^11^ highlights a connection between GSK3*β* and FRAT2, based on the crystal structure of the complex formed by GSK3*β* and FRAT1, a close paralog of FRAT2. But it shows no connection between them and the exportin XPO1. The latter is only linked to erbB2, which shows no link to either GSK3*β* or FRAT2. Surprisingly, a NES motif links erbB2 with XPO1 (dashed pink line in Fig. 3C) but it seems unlikely as all disorder prediction scores are <0.15 and it lies within the extracellular and structured right-helix *β*-sheet of the Receptor L domain (PF01030). The older server Interactome3D^9^ failed to link GSK3*β* to any partner in absence of detected complexes involving this protein (Fig. 3B). It seems, here, that the segment of FRAT1 bound to GSK3*β* is too small to be taken into account for modeling FRAT2-GSK3*β* interaction. The well-known STRING-db^7^ server provides many links within the submitted 14-member networks but none connecting directly GSK3*β* or FRAT2 with XPO1 (Fig. 3A). While, PrePPI ^18^ indicates an experimental connection (from BioGRID) between FRAT2 and XPO1, no motif is highlighted and no prediction score is provided for that interaction. Hence, those servers could not provide a clear explanation for the GSK3*β* translocation. Similarly, Uniprot does not described the NES motif in neither FRAT1 nor FRAT2. However, it does provide a link to ComplexPortal describing a GSK3*β*-FRAT2 complex involved in nuclear export (entry CPX-462) but without explicit connection with the exportin XPO1. On the contrary, using SLiMAn, we straightforward produce a convincing atomic resolution model, potentially explaining the GSK3*β* shuttling from nucleus to cytoplasm, which indeed results from a combination of the domain-domain interaction of GSK3*β* with FRAT2, and a SLiM-domain recognition of the FRAT2 NES motif by the exportin XPO1.

**Figure 3:**
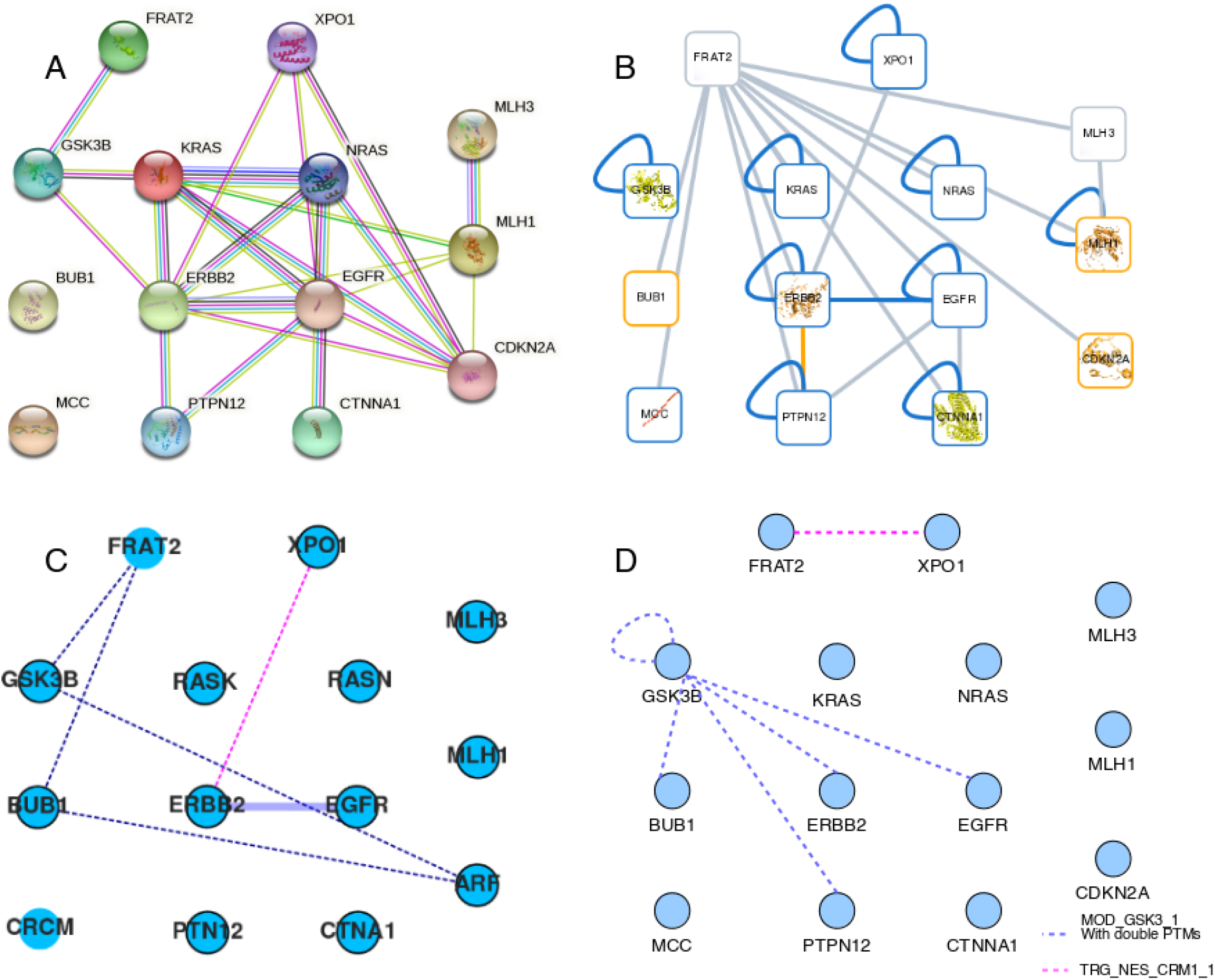
Networks of the 14 BioGRID partners for FRAT2 generated by **A)** STRING-db,^7^ **B)** Interactome3D,^9^ **C)** Proteo3DNet^11^ and **D)** SLiMAn (this work). **A)** Output screen-shot from String-db. Links are colored using default settings from the server. **B)** Output screenshot from Interactome3D. Links are colored using default settings the server. **C)** Links extracted from Proteo3DNet. Thick lines correspond to domain-domain interactions, while dashed lines highlight ELM-based connections in pink (erbB2-XPO1) or blue (protein phosphorylation), respectively. **D)** Links extracted from SLiMAn, at high and low confidence, are highlighted in pink (FRAT2-XPO1) and blue (GSK3*β* phosphorylation) dashed lines, respectively.

While the interaction between FRAT2 and GSK3*β* are not suggested by other web servers (Fig. 3), the latter provide information for other putative interactions. Indeed, the other tools suggests a core network around erbB2/EGFR. This is not highlighted by SLiMAn and likely relies on interactions through folded regions. For example, the dimerization of erbB2 and EGFR involves a well characterized transmembrane domain interaction (see PDB2KS1). They also display an interaction between XPO1 and erbB2, based on a co-immunoprecipitation assay.^31^ Additionally, various connections with GSK3*β* are suggested especially by Proteo3Dnet although this mainly relies on phosphorylation by various other protein-kinases (see above). Nevertheless, this prompted us to dig into SLiMAn results. To show motifs for phosphorylation by GSK3*β* (MOD GSK3 1), we set the E-value threshold above 0.027 and all IUpred scores set to ≥ 0.5. Changing this parameter increases the number of possible interactions to 63 (from 12). Among these putative interactions, SLiMAn listed many GSK3*β* motifs among which 10 are doubly phosphorylated sites according to PhosphoSitePlus®. Accordingly, this additional analysis suggests that 5 of the 14 partners of FRAT2 are substrates of GSK3*β*.

In conclusion, all these tools are complementary, and SLiMAn appears optimal for SLiM-based interactions analysis.

### Pragmin interactome

Hereafter, we described a follow-up study of the interactome of Pragmin. ^27^ We previously described automatic 3D modeling of several complexes contained in this interactome. We resume and extend the automatic analysis of ELM/Pfam matches here using SLiMAn. This new search revealed new interactions within the submitted list of 62 proteins.

As this interactome was not deposited in BioGRID, we can used BioGRID information as a first-step validation while adding information from putative ELM motif/Pfam domain matches to highlight putative modes of interaction. From previous works on pragmin and its close homlogues PEAK1 and PEAK3, two principal interactions through SLiMs were known. This includes the recognition of a specific EPI(Y)A motif (in which the tyrosine is phosphorylated) in pragmin by the SH2 domain of Csk and, by similarity, the binding of CrkL to a poly-proline motif also present in the long N-terminal intrinsically disordered region (960 aa) of pragmin. By default, SLiMAn suggests more direct partners for Pragmin including SH2 and SH3 containing proteins (e.g.: Grb2 or Crk) as well as the phosphatase PP2BA (see below). But we resume our search following a hierarchical strategy starting from the most likely interactions or motifs.

First, we filtered the search for true instances from ELM, highlighted two interactions through a phosphorylated SH2 motif (LIG SH2 CRK) in Paxillin and Crk, both possibly recognized by Crk (http://sliman.cbs.cnrs.fr/pragmin/Instances). These interactions are sustained by two low-throughput and one high-throughput data in BioGRID, respectively. Interestingly these two proteins are well-known partners of other members of this interactome, although the corresponding interactions are not yet appearing here. Then, we turned to evaluate the possible pairs involving a SLiMs predicted to be in a strict disordered state (all 5 IUpred scores ≥ 0.5) one ELM class at a time. We noticed no cleavage (CLV) site nor degron (DEG) motif corroborated by a BioGRID match. One targeting motif (TRG ER diLys 1; E-value = 2.7 e-5) linked two proteins: the proline hydroxylase P3H1 and Kinesin-like protein KI21B. Motifs from the LIG and DOC classes brought 27 and 7 pairs, respectively. SH2, WW and SH3 motifs were connecting Crk and CrkL to Grb2 while paxillin can be linked to Csk, Crk, Grb2, CrkL and Stxb4. These putative interactions are supported by various experimental data listed in BioGRID. In turn, it is predicted to provide many domains (mainly SH2 and SH3) and additional motifs for further interactions with other partners. Accordingly, this would represent a macromolecular assemblage attached to pragmin through at least Csk and CrkL, not envisioned before.

In addition, SLiMAn highlighted a possible dimerization (or internal looping) of the Abl inhibitor 1 through its SH3 motif and its own SH3 domain as well as a connection between the protein kinase DyrK1a with the 14-3-3 protein *η*. PhosphoSitePlus® confirmed this site to be phosphorylated and the literature suggested it would correspond to an inhibited form of the DyrK1A. So far, those two proteins were not connected to pragmin nor to its close interactants. The DOC class comprises 7 docking motifs specific for protein-kinases not present in the pragmin interactome. As noted above, the PFAM classification pools most of the protein-kinases in the PF00069 superfamily. So, improper pairing is likely to occur. Accordingly, the proposed pairings of AAKB1 and AAKG2 with AAPK1 or DyrK1a with itself and KCC2D were deemed inconsistent. The BioGRID links correspond to the association of two regulatory subunits, AAKB1 and AAKG2, with the catalytic subunit AAPK1 to form a functional AMPK (in agreement with Proteo3DNet ^10^) or to some kind of dimerization for DyrK1a. The interaction of the latter with the protein-kinase KCC2D would require further investigation. In the MOD class, 22 pairings are reported by SLiMAn but again with a lack of specificity for the domain definition (PF00069) for most of the phosphorylation sites but in two cases. The AMPK regulatory subunit AAKG2 harbors an accessible motif for phosphorylation by DyrK1a while Abi1 would be phosphorylated by AMPK. In both cases, BioGRID enlisted an experimental interaction. The corresponding results can be viewed on our web server (http://sliman.cbs.cnrs.fr/pragmin/StrictDisorder). Noteworthy, without constraint on BioGRID connections, 310 pairings are predicted with only one more linked to DyrK1a and three corresponding to the protein-kinase Plk1. However, only two of these motifs are supported by the data in PhosphoSitePlus® and would correspond to phosphorylation of apoptosis-inducing factor mitochondria-associated 1, AIFM1, and the chaperone, HS105, by Plk1.

Lowering further the stringency (IUpred scores kept to ≥ 0.5 but LongDomain set to 0 and ANCHOR2 set to 0.3) of our search revealed other pairings validated by BioGRID. The various ELM classes TRG, DEG, LIG, DOC and MOD include one, two, four, zero and two additional pairings, respectively. This brings new partners such as the coiled-coil containing protein CCD33 and the chaperone BIP (with Stxb4 through a WW and a PDZ motif, respectively) or Dcaf7, the DDB1- and CUL4-associated factor 7 (with Plk1 through a DEG motif and with DyrK1a with a TRG motif). If data from BioGRID are no longer taken into account, SLiMAn connects the tankyrases to PDIP2 and A2AP1 while PP2BA is predicted to recognize its specific PP2B motifs in pragmin, paxillin and others (AP2A1, WDCP, DyrK1a, KI21B) and a tankyrase (TNKS1). Noteworthy, a recent proteomics study^32^ using the tankyrases as baits identify A2PA1 and two other members of the pragmin interactome (Kelch-like protein 7 or KLHL7 and the Nck-associated protein 1, NGAP). This highlights that pinpointing the most likely interactions through ELM motifs/Pfam domain pairing allow one to focus efficiently the research of true interactants in the litterature. The corresponding results can be viewed on our web server (http://sliman.cbs.cnrs.fr/pragmin/Intermediate).

Using default parameters (IUPRED set to 0.3 but LongDomain, ShortDomain and ANCHOR2 set to 0, 1 and 0.5, respecctively), SLiMAn highlights 81 possible interactions (38 without phosphorylation sites) out of 26894 pairs matching one ELM motif (out of 1718 detected motifs) and one Pfam (out of 30) domain annotations. It includes motifs from different ELM classes (O CLV, 0 DEG, 1 TRG, 47 LIG among which 12 with required phosphorylation that are actually observed, 6 DOC and MOD motifs). Clearly, this highlights the power of filtering out a huge amount of unlikely pairings while the above association readily provide clues to explain the presence of various proteins within the interactome of pragmin. Lowering further the stringency and careful scrutiny led us to identify more putative interactions among the 62 proteins of the pragmin interactome, which leave only 9 proteins singletons.

This work also recapitulates part of our former manual analysis showing a cluster of interactants around pragmin including Csk, Crk, Crkl and Grb2 as previously described. Obviously, more analysis and experimental studies are necessary to confirm those hypothesis but a clearer picture of this interactome is emerging with many partners shown as likely involved in dedicated interactions in the surrounding of pragmin. Some interactions are already linked to functional outputs linked to cancer and could explain the oncogenic behaviour of pragmin.

As in the above two examples, most of the interactions were validated through comparative modeling of the corresponding complexes. However, they are awaiting further experimental validations but SLiMAn clearly points to hot-spots to guide these experimentations.

## Conclusion

This manuscript describes a new tool, named SLiMAn and developed to interrogate simultaneously the databases of short linear motif ELM and of conserved domains Pfam, through an user-friendly interface. Users should provide, as input, a list of putatively interacting proteins (such as an output from an interactomics study) to SLiMAn server, to quickly highlight the ELM motifs present in those sequences and paired with Pfam domains present within the same set of proteins. Importantly usual search parameters in the field of SLiMs analysis (E-value, disorder, …) can be adjusted to search for alternative pairings. When possible, sequence to structure alignments with related 3D templates are provided to enable comparative modeling the binary complex. This help highlighting potential physical interactions for a short list of proteins. Motifs often correspond to weak sequence matches in more or less disordered region making them tricky to find. Fuzzy logic is necessary to attempt to filter in true positives. Focusing on pairs of ELM motifs and Pfam domains allow users to better scrutinize a focused set of potential interactors. We provided three distinct examples of application. In one case, interrogating the BioGRID interactome of FRAT2 with SLiMAn revealed its potential physical interactions with XPO1/Exportin. This link explains the well-known shuttling of GSK3*β* (Bechard, et al., 2012) and represents a potential target against some cancers. Such an example illustrates the benefit of SLiMAn to rapidly reveal potentially direct interactions in a sub-proteome.

In conclusion, SLiMAn shall nicely complement current tools and databases – such as STRING-db,^7^ BioGRID,^8^ or Proteo3Dnet^11^ – by connecting efficiently proteomics data with ELM motifs^14^ and Pfam domains^15^ . To our knowledge, there is currently no tool or web server equivalent to SLiMAn. Accordingly, SLiMAn represents an interesting new tool in the field of protein-protein interactions to rapidly and precisely pinpoints potential SLiM-based contacts among a focused list of proteins.

## Acknowledgement

The authors thanks Jean-Luc Ponsfor careful reading of the manuscript and hardware survey. We are grateful for the comments from editors and referees.

## Supporting Information Available

The following supporting information is available free of charge at ACS website http://pubs.acs.org

Supplementary Table 1. Headers description of the hit predictions results

Supplementary Table 2. Details on the filtering panel : parameters and values

Supplementary Table 3. Content of the summary tables after filtering

## TOC Graphic

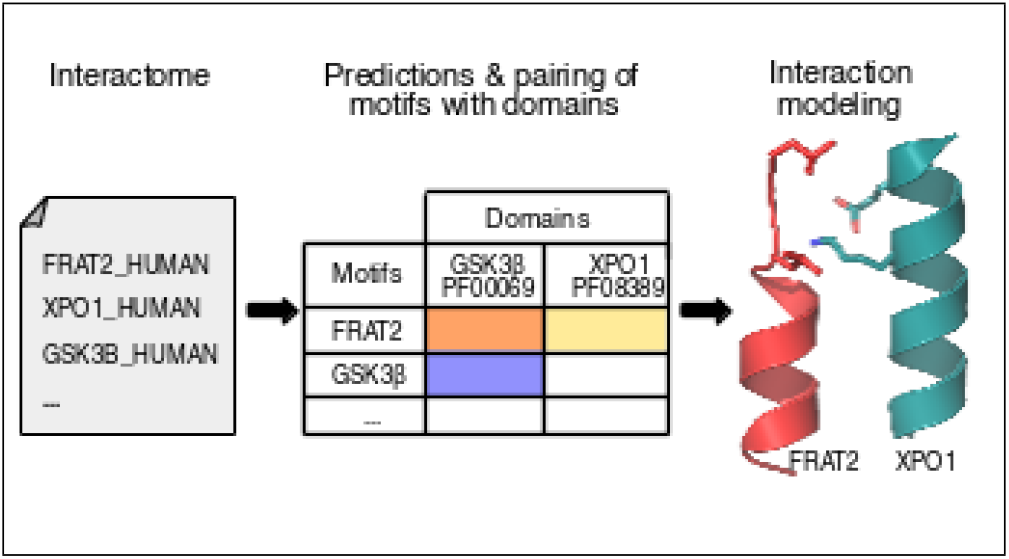

