## Supplementary material for "SLiMAn: an integrative web server for exploring short linear motif-mediated interactions in interactomes": Supplmental Information


### Supporting Information Available

The following supporting information is available free of charge at ACS website `http://pubs.acs.org`

Supplementary Table 1. Headers description of the hit predictions results

Supplementary Table 2. Details on the filtering panel : parameters and values

Supplementary Table 3. Content of the summary tables after filtering

[†]Short Linear Motifs Analysis

Table 1: Hit prediction file headers descriptions

| Name | Short description | Values |
|---|---|---|
| ELM_CLASS | Class of matched ELM motif | Class Name |
| MATCHED_MOTIF | Motif extracted from the sequence | Amino acids sequence |
| MOTIF_UNICODE | Uniprot acc. of the protein | Uniprot acc. |
| MOTIF_STARTS | Start of the motif in the protein sequence | integer |
| MOTIF_ENDS | End of the motif in the protein sequence | integer |
| ASSOCIATED_PFAM | Pfam domain identifier of the motif associated domain | PfamID |
| MATCHED_PFAM_UNICODE | Uniprot acc. of the protein containing the Pfam domain | Uniprot accession |
| EXPERIMENTAL_EVIDENCE | ELM valid instance | $\{0, 1\}$ |
| ELM_PROBA | Motif E-value | $[7.24 \times 10^{-16}, 0.018]$ |
| STRICT_DISORDER | All IUpred scores above 0.5 | $\{0, 1\}$ |
| SHORT_IUpred_AVG_SCORE | IUpred short window average score | [0, 1] |
| LONG_IUpred_AVG_SCORE | IUpred long window average score | [0, 1] |
| SHORT_IUpred_GLOB_DOM | IUpred short window globular domain average score | [0, 1] |
| LONG_IUpred_GLOB_DOM | IUpred long window globular domain average score | [0, 1] |
| ANCHOR2_AVG_SCORE | IUpred Anchor2 average score | [0, 1] |
| BioGRID_INTERACTIONS | BioGRID low throughput interactions count | $\geq 0$ |
| BioGRID_EXFILTERED | BioGRID high throughput interactions count | $\geq 0$ |

Table 2: Parameter panel

| Main parameter | Parameter | Short description | Value range | Default value |
|---|---|---|---|---|
| ELM | Motif E-value | Motif E-value | $[7.24 \times 10^{-16}, 0.018]$ | 0.005 |
| | Verified Instances | 1 if predicted interaction is part of the ELM valid instances else 0 | {0, 1} | 0 |
| | Class types | Modification sites (MOD) | {0, 1} | 1 |
| | | Docking sites (DOC) | {0, 1} | 1 |
| | | Ligand binding sites (LIG) | {0, 1} | 1 |
| | | Degradation sites (DEG) | {0, 1} | 1 |
| | | Targeting sites (TRG) | {0, 1} | 1 |
| | | Cleavage sites (CLV) | {0, 1} | 1 |
| IUpred | Strict Disorder | 1 if all IUpred scores $\geq 0.5$ else 0 | {0, 1} | 0 |
| | Short avg. score | Lb.[a] IUpred short window average score | [0, 1] | 0.3 |
| | Long avg. score | Lb.[a] IUpred long window average score | [0, 1] | 0.3 |
| | Short Glob.Dom. avg. score | Lb.[a] IUpred short window glob. dom.[b] average score | [0, 1] | 1 |
| | Long Glob.Dom. avg. score | Lb.[a] IUpred long window glob. dom.[b] average score | [0, 1] | 0 |
| | Anchor2 avg. score | Lb.[a] IUpred Anchor2 average score | [0, 1] | 0.5 |
| BioGRID | Low Throughput | Lb.[a] low throughput BioGRID interactions | $\geq 0$ | 0 |
| | High Throughput | Lb.[a] high throughput BioGRID interactions | $\geq 0$ | 0 |
| | Total count | Lb.[a] low and high BioGRID interactions | $\geq 0$ | 1 |
| SLiMAn | Confidence level | Lb.[a] the confidence level | {1, 2, 3, 4} | 1 |
| | Available template | Lb.[a] nb. of available templates for the alignment | $\geq 0$ | 0 |
| | Validated Models | Lb.[a] nb. of validated models | $\geq 0$ | 0 |
| | Sorting Algorithm | Alphabetically (Alphabetic) | {0, 1} | 0 |
| | | User order (Input Order) | {0, 1} | 1 |
| | | Match similarities (Clusters) | {0, 1} | 0 |

[a] Lower boundary of; [b] Globual Domain prediction

Table 3: Summary tables

| | | | |
|---|---|---|---|
| Proteins | | | |
| | Filtered IN | Filtered OUT | Not found |
| ELM | Nb. prot. for which at least one motif is displayed | Nb. prot. for which no motif is displayed | Nb. prot. containing no motif |
| Pfam | Nb. prot. for which at least one domain is displayed | Nb. prot. for which no domain is displayed | Nb. prot. containing no domain |
| Entries | | | |
| | Filtered IN | Filtered OUT | Sum |
| ELM | Nb. of unique ELM classes displayed | Nb. of ELM classes filtered out | All ELM class found for the run |
| Pfam | Nb. of unique domains displayed | Nb. of unique domains filtered out | All Pfam domains found for the run |
| Total Interactions | | | |
| | Filtered IN | Filtered OUT | Sum |
| ELM | Sum of displayed ELM classes | Sum of filtered out ELM classes | Sum of ELM classes found for the run |
| Pfam | Sum of displayed domains | Sum of filtered out domains | Sum of domains found for the run |
| SLiMAn | Sum of displayed hits | Sum of filtered out hits | Sum of all SLiMAn hits found for the run |
